# Prohibitin 2 is a Key Regulator of T Cell Proliferation and Effector Functions

**DOI:** 10.1101/2024.12.13.628303

**Authors:** Johannes F. Vogt, Carsten Merkwirth, Petra Adams-Quack, Elena Zurkowski, Thomas Michna, Assel Nurbekova, Ute Distler, Hajime Yurugi, Sonja Reißig, F. Thomas Wunderlich, Krishnaraj Rajalingam, Stefan Tenzer, Axel Methner, Thomas Langer, Ari Waisman, Nadine Hövelmeyer

## Abstract

Prohibitin 2 (PHB2) is a highly conserved protein with essential roles in cell homeostasis and survival across different cell types. Previous studies have shown that the deletion of PHB2 results in an arrest in proliferation due to impaired mitochondrial function regulated by the dynamin-like GTPase OPA1. The function of PHB2 in immune cells remains unclear, however, some studies suggest that PHB2 plays a role in the cell membranes of B and T cells. In order to elucidate the role of PHB2 in immune cells, we generated PHB2-deficient T cells. Our findings reveal a pivotal role for PHB2 in the proliferation and differentiation of T cells. PHB2 deficiency inhibits T cell proliferation by inducing a cell cycle arrest at the G1 to S phase, thereby preventing the differentiation into effector T cells. Furthermore, in contrast to previous reports, T cells lacking PHB2 are more resistant to apoptosis. Metabolic analysis reveals that PHB2-deficient T cells fail to upregulate their glycolysis and oxidative phosphorylation upon activation, thus rendering them incapable to proliferate.

## Introduction

Prohibitin 1 (PHB1) and prohibitin 2 (PHB2) are highly conserved proteins, ubiquitously expressed in essentially all cell types, ranging from murine embryonic fibroblasts, intestinal epithelial cells, β-cells to neurons^1–4^. PHB1 and PHB2 are interdependent on the protein level, and loss of one of them simultaneously leads to loss of the other^1^. Prohibitins exert and regulate a diverse range of cellular functions, including apoptosis, energy metabolism, proliferation, and senescence, based on their cellular location, either at the plasma membrane, mitochondria, or the nucleus^5, 6^.In mitochondria, both proteins assemble at the inner mitochondrial membrane and form a supra-macromolecular structure that regulates the mitochondrial genome, modulates mitochondrial dynamics, morphology, biogenesis, and the mitochondrial intrinsic apoptotic pathway^7, 8^.

While mitochondrial Phbs are always present in all cell types studied so far, a considerable proportion of PHB1 and PHB2 were shown to be expressed on the surface of T cell receptor (TCR) activated T cells in mice and humans^6, 9^. Specifically, PHB1 and PHB2 are highly expressed on the surface of Th17 cells, contributing to the differentiation into these effector cells^10^. Engagement of surface-expressed PHB1/2 with Vi polysaccharide leads to the inhibition of the CRAF-MAPK signaling cascade that controls the plasticity of Th17 cells, although this interaction has no effect on the proliferative capacity of T cells^10^. The knockdown of prohibitins in Kit255 cells, a T cell leukemia cell line, led to a reduced mitochondrial membrane potential^9^. This suggests that prohibitins on the cell surface of activated T cells are involved in TCR-mediated signaling^6^. Although these *in vitro* studies strongly indicate a role of prohibitins in T cell function, *in vivo* evidence is still lacking.

In mice, it was shown that loss of the PHB complex during development and mice brain-specific deletion of *Phb2*, using Nestin-Cre, results in embryonic lethality^11^. Postnatal knockout of PHB2 in the forebrain using *CaMKIIα*- driven cre expression leads to neuronal loss and death at around 17 weeks of age^4^. To determine the function of PHB2 in T cells, we examined mice with a T cell-specific PHB2-deficiency and found that PHB2 is essential for T cell homeostasis, proliferation, and T cell differentiation. At steady state, the T cell numbers of mice lacking PHB2 were significantly reduced in all secondary lymphoid organs tested. Even though PHB2-deficient T cells still show typical early TCR activation signs, such as increased expression of CD25 and CD69, these naive T cells are unable to differentiate into effector T cells and to secrete effector cytokines upon activation. Furthermore, T cells lacking PHB2 exhibit a significant proliferative defect resulting from cell cycle arrest at G1 to S-phase. This is due to a defect in initiating DNA replication following TCR stimulation, which is accompanied by a significantly reduced expression of numerous proteins crucial for T cell proliferation. Thus, our study highlights the significance of PHB2 for T cell functions *in vivo*.

## Results

### PHB2 is essential for T cell homeostasis *in vivo*

To study the role of PHB2 in T cells *in vivo*, we crossed conditional *Phb2^F/F^* ^1^ mice with CD4-Cre^12^ mice, resulting in the deletion of PHB2 in all T cells from the CD4^+^CD8^+^ double-positive progenitor state during thymic development. For brevity, these mice are referred to as *Phb2^TKO^*.

*Phb2^TKO^* mice showed no macroscopic abnormalities and developed comparable to littermate control mice (data not shown). Even though CD4-Cre expression starts in the double-positive state of T cell development in the thymus we did not detect any changes in thymocyte cell numbers or subpopulations by flow cytometric analysis of *Phb2^TKO^* mice compared to littermate controls (Fig. S1A, B).

The total cell counts of the spleen, node (LN) and mesenteric lymph nodes (mLN) were also not significantly different between the different genotypes (Fig. S1A). However, the percentage as well as the total cell count of TCRβ^+^ T cells in the spleen was reduced by half and accompanied by a significant increase in the percentage but not in total cell count of CD19^+^ B cell numbers (Fig. 1A). The ratio of splenic CD4^+^ and CD8^+^ T cells remained unchanged while there was a significant reduction in the total cell count compared to littermate controls (Fig. 1B). Similar effects were observed for both LN and mLN (Fig S1C). Furthermore, when CD4^+^ T cells were analyzed for the expression of activation markers, we again observed a significant reduction when subdividing the CD4^+^ T cells into naïve (CD62L^high^CD44^low^) and memory/effector T cells (CD62L^low^CD44^high^) (Fig.1C). The total cell counts of naïve and central memory CD8^+^ T cells was also significantly reduced, whereas the number of effector CD8^+^ T cells remained unchanged (Fig. 1D.) Another T cell subset that is affected by CD4-cre are Foxp3^+^ regulatory T (Treg) cells. Flow cytometric analysis of splenic Treg cells again showed a significant reduction in this cell population in percentage as well as total cell count compared to controls (Fig. 1E). To understand why thymic T cells were unaffected by the loss of PHB2 we tested CD4-cre mediated deletion efficiency of PHB2 by Western blot analysis of MACS purified CD4^+^ thymic T cells. Even though CD4-Cre expression begins in the double-positive state of T cell development, PHB2 was still present at equal levels in *Phb2^TKO^* thymocytes compared to control cells (Fig. S1D), suggesting that the half-life for PHB2 is long and deletion manifests only in the periphery.

**Figure 1:**
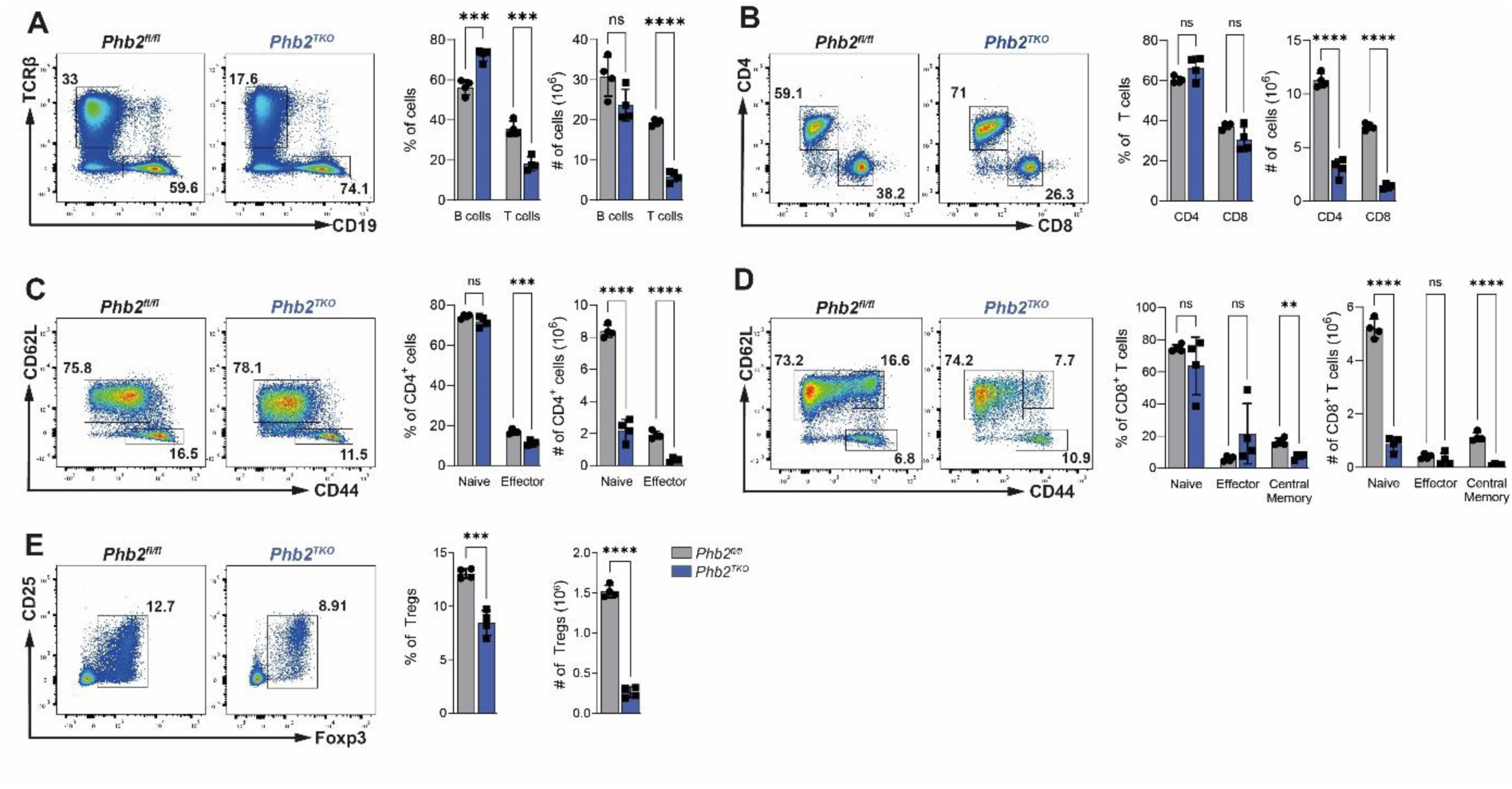
Prohibitin 2 is essential for T cell homeostasis in vivo. A: Flow cytometric (left) and statistical analysis (right) of live B cells (CD19 ^+^) and T cells (TCRβ^+^) B: Flow cytometric (left) and statistical analysis (right) of live CD4^+^ and CD8 ^+^ T cells C: Flow cytometric (left) and statistical analysis (right) of live CD4 naïve (CD62L^+^ CD44^lo^ ^w^) and CD4 effector ( CD62L^-^ CD 44 ^h igh^) T cells D: Flow cytometric (left) and statistical analysis (right) of live CD8+ naïve (CD62L^+^ CD44^lo^ ^w^), effector CD8+ ( CD62L^-^ CD44 ^h igh^), and central memory CD8+ ( CD62L^+^ CD44^h^ ^i gh^) T cells E: Flow cytometric (left) and statistical analysis (right) of live Treg cells (Foxp3 ^+^) Data is representative of three independent experiments with n=4. Bar graphs show means +/- SDs and single values. Statistical significance was calculated using unpaired two-tailed t-test with Holm-Šidák correction for multiple comparison. **p≤0.01 ***p≤0.001, **** p≤0.0001.

Western blot analysis of FACS sorted naïve CD4^+^ and CD8^+^ T cells revealed a complete deletion of PHB2 protein (Fig.S1E). In contrast, the CD4^+^ and CD8^+^ effector T cells of *Phb2^TKO^* mice contained some residual PHB2 protein (Figure S1E).

Since CD4-cre also targets Treg cells, we sorted CD4^+^CD25^+^ T cells for Western blot analysis to check deletion efficiency in this T cell population. As expected, this analysis revealed an almost complete loss of PHB2 protein in Treg cells (Fig. S1F). Furthermore, as reported previously^1^, the deletion of *PHB2* also led to the loss of PHB1 on protein level (Fig. S1G).

In summary, the deletion of PHB2 in T cells results in a dramatic decrease of total T cells at steady state in peripheral lymphoid organs. These results show that the loss of PHB2 in T cells prevents appropriate T cell expansion in the periphery, while T cell development appears to be normal.

### Prohibitin 2 is essential for *in vivo* effector T cell function

To assess the function of PHB2-deficient T cells *in vivo*, we induced experimental autoimmune encephalomyelitis (EAE). EAE is a mouse model of multiple sclerosis and is highly dependent on T cell effector functions. This autoimmune reaction is activated by injecting the MOG peptide p35-55 emulsified in CFA, which activates autoimmune T cells and promotes their migration into the central nervous system^13^. This process leads to CD4^+^ T cell-mediated destruction of neuronal myelin sheaths in the central nervous system.

We found that mice lacking PHB2-specifically in T cells were completely resistant to EAE compared to control animals (Fig. 2A, B). In order to rule out a paracrine effect, we generated bone marrow (BM) chimeras. We isolated BM cells from Thy1.1 control and Thy1.2 *Phb2^TKO^* mice, and these cells were injected (in a ratio of 1:1) into Ly5.1 mice that were previously irradiated. ten weeks later, EAE was induced in these animals. The disease progressed as expected, and the mice were analyzed at the peak of the disease on day 15 by flow cytometry (Fig. 2C). This analysis revealed that PHB2-deficient CD4^+^ T cells were absent in the central nervous system (CNS) of sick mice, which only contained Thy1.1 WT T cells (Fig. 2D). Furthermore, the MOG recall-assay, an assay for MOG specific auto-immune T cells, showed a significant reduction in CNS infiltration by MOG-specific PHB2-deficient CD4^+^ T cells (Fig. 2E).

**Figure 2:**
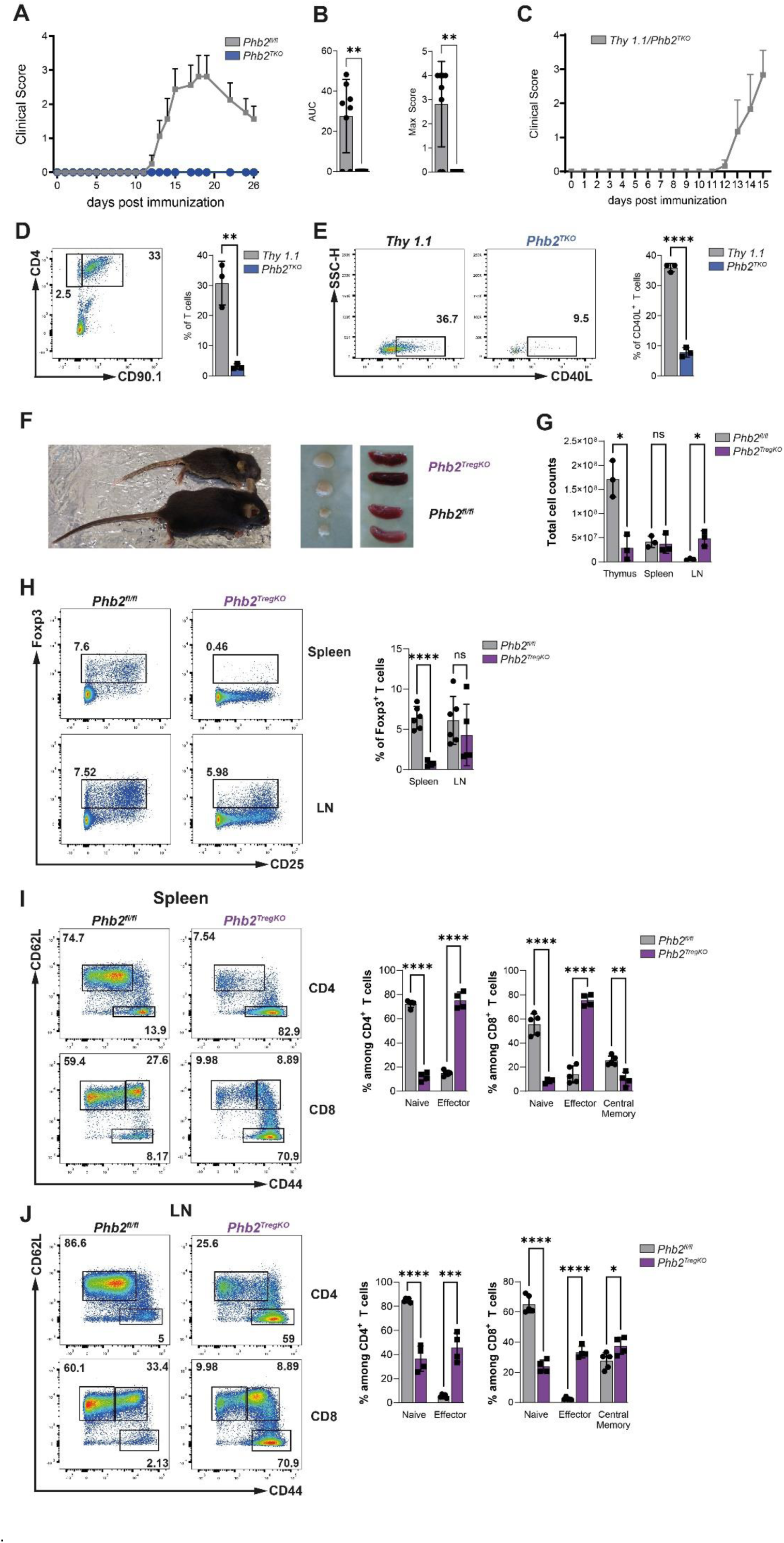
PHB2 is essential for T cell effector function. (A) Clinical signs of EAE are shown as mean clinical disease scores ± SD. (n=5-7) (B) Area Under the Curve (AUC) and maximum clinical scores of (A). n=7-8, Mann-Whitney test ±SD (C) Clinical signs of EAE of bone marrow chimera of *Thy1.1* and *Phb2^TKO^* mice. n=3 (D) Flow Cytometric (left) and statistical analysis (right) of CD4^+^ CD90^+^ T cells from the CNS of mice from C (E) Flow Cytometric (left) and statistical analysis (right) of CD40L^+^ T cells from the CNS of mice from C (F) Representative picture of 29 days old control and *Phb2^TregKO^* mice and their spleen and inguinal lymph nodes (G) Total cell count of thymus, spleen and LN of 29 days old control and *Phb2^TregKO^* mice. n=3 (H) Flow Cytometric (left) and statistical analysis (right) of Tregs (Foxp3+). (n=4-6) (I,J) Flow Cytometric (left) and statistical analysis (right) CD4^+^ naïve (CD62L^+^ CD44^low^) and CD4^+^ effector (CD62L^-^ CD44^high^) T cells and naïve CD8^+^ (CD62L+ CD44^low^), CD8^+^ effector (CD62L^-^ CD44^high^), and central memory CD8^+^ (CD62L^+^ CD44^high^) T cells in Spleen (**I**) and LN (**J**) Data is representative of three (A) one (C-E) or pooled from two (G-J) independent experiments. Bar graphs show means +/- SDs and single values (D,E,G,H,I,J) Statistical significance was calculated using unpaired two-tailed t-test with Holm-Šidák correction for multiple comparison. *p<0.05, **p≤0.01, ***p≤0.001, **** p≤0.0001.

These results show that T cells lacking PHB2 are incapable of responding to the MOG peptide used for the immunization. Given that *Phb2^TKO^* mice are phenotypically normal, yet their T cells are incapable of mounting an immune response, we decided to cross *PHB2^fl/fl^* mice with Foxp3-Cre mice to investigate potential defects in PHB2-deficient regulatory T cells.

*Phb2^TregKO^* mice exhibited clear signs of autoinflammation and had to be sacrificed between 22-32 days after birth due to severe signs of inflammation. Macroscopically, *Phb2^TregKO^* mice were smaller than control mice and displayed scaly skin, crusting of the eyelids, ears and tail, signs of a scurfy phenotype (Fig. 2F)^14^. These mice showed enlarged lymph nodes (LN) while the spleens were similar in size compared to those of control mice (Fig. 2F). Consistent with its size, the total cell counts of the spleen remained unchanged when comparing *Phb2^TregKO^* with control mice. However, the peripheral LN of *Phb2^TregKO^* mice displayed a marked increase in total cell count (Fig. 2G). A detailed analysis of the thymus revealed a dramatic reduction in the total cell count (Fig. 2G).

Flow cytometric analysis revealed a significant reduction in the percentage of Treg cells in the spleen, while the LN remained unchanged (Fig. 2H). Additionally, analysis of effector T cells showed a dramatic increase in CD4^+^ and CD8^+^ effector T cells in both the spleen and LN of *Phb2^TregKO^* mice compared to control mice (Fig. 2I, J). Together these results resemble the scurfy phenotype, an X-linked recessive mutation, that leads to a loss of Treg function and subsequent fatal immune dysfunction^14^. These findings collectively underscore the critical importance of PHB2 for the functionality of T reg cells.

### PHB2 regulates proteins that are crucial for T cell proliferation

In order to obtain an unbiased view of the effect of PHB2 deficiency on T cells, we performed a comparative proteomic analysis of MACS purified splenic naïve CD4^+^ T cells from *Phb2^TKO^* and control mice at steady state and after α-CD3/α-CD28 stimulation for 24 hrs (Fig. 3).

**Figure 3:**
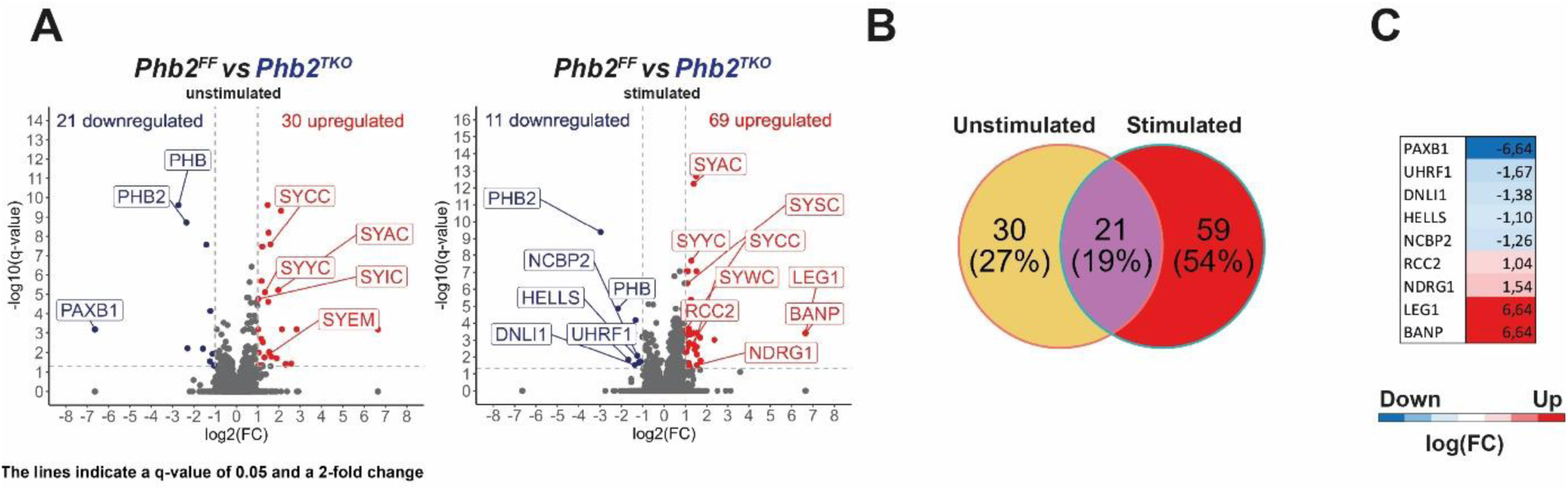
PHB2 regulates proteins that are essential for proliferation. (A) Volcano plots of mass spectrometric analysis of unstimulated (left) and α- CD3/α-CD28 stimulated (right) naïve CD4 ^+^ T cells (B) Venn diagram of significantly regulated peptides in unstimulated and stimulated naïve CD4 ^+^ T cells (C) Manually annotated list of significantly regulated peptides with a negative effect on proliferation

At steady state, T cells from *Phb2^TKO^* mice were significantly different from T cells from control mice. In unstimulated PHB2 deficient naïve CD4^+^ T cells, 21 peptides were significantly downregulated and 30 peptides upregulated compared to naïve control T cells (Fig. 3A, left).

After 24h of α-CD3/α-CD28 TCR stimulation, eleven peptides were significantly downregulated and 69 peptides were upregulated in PHB2-deficient T cells compared to control T cells (Fig. 3A, right). Of these, 21 peptides were significantly altered under both unstimulated and stimulated conditions (Fig. 3B, Fig. S3A). Surprisingly, the changes in expression levels of these 21 peptides were identical under both conditions, suggesting that the stimulation itself did not affect the expression of these peptides (Fig. S3A). This analysis also revealed the loss of both PHB2 and PHB1 in the transgenic T cells compared to control mice (Fig S3A). At steady state and after stimulation, annexin A2 (Anxa2), a protein that has already been reported to be directly associated with PHB1 and PHB2^15^, was upregulated in PHB2- deficient T cells compared to control cells (Fig. S3A). After stimulation with α-CD3/α-CD28 the ATP-dependent Zinc metalloprotease YME1L1 was downregulated in PHB2-deficient T cells. YME1L1, in conjunction with OMA1, cleaves OPA-1, resulting in a disbalance in the S-OPA1/L-OPA1 ratio. This imbalance is responsible for the fragmented mitochondrial phenotype observed under PHB2 deficiency^116^. YME1L1 downregulation in PHB2-deficient T cells in comparison to control T cells, suggests a potential compensatory mechanism (Fig. S3A).

This data set was further analyzed using the Ingenuity Pathway Analysis (IPA). This pathway analysis revealed no significant regulated pathways related to proliferation, metabolism or T cells. However, tRNA charging was significantly upregulated in PHB2-deficient T cells under steady state and after stimulation in comparison to control T cells (Fig. S3B). This pathway consists of tRNA synthetases, ligases that mediate the loading of tRNA with amino acids^17^. Furthermore, IPAs functional analysis revealed that PHB2-deficient T cells exhibit an increase in the expression of peptides associated with T cell activation and T cell development when compared to control cells (Fig. S3C).

Next, we manually annotated our data set and found a set of peptides associated with cell proliferation. We found nine differentially expressed peptides that are associated with proliferation and cell cycle progression (Fig. 3C), five of those with indication for a direct role in T cell proliferation (Supple. Table 1). All these peptides were regulated in a way that they would negatively affect the proliferation of T cells. Out of these proteins HELLS, a helicase that establishes DNA methylations patterns, has already been reported to affect peripheral lymphocyte homeostasis and T cell proliferation^18^ but was so far not shown to be associated with prohibitins.

Collectively, these data suggest a pro-proliferative role of PHB2 in T cells, as indicated by the downregulation of proteins essential for proliferation.

### PHB2 is essential for proper T cell function

To determine if the downregulation of proliferation-associated proteins observed in the proteomics data set from PHB2-deficient T cells leads to a proliferative defect and a subsequent reduction in all peripheral T cell subsets, we labeled naïve CD4^+^ T cells with a violet cell tracer (VCT) and activated via the TCR. In addition, we added cytokines cocktails to induce their differentiation to Th1, Th17, or Treg cells. We detected a profound proliferative defect of PHB2-deficient T cells independent of the stimulation cocktails used (Fig. 4A). To better characterize Th cell effector function we investigated cytokine secretion by CD4^+^ T cells from *Phb2^TKO^* and control mice. Differentiation of naïve CD4^+^ T cells into Th17 and iTregs for four days revealed a significant reduction of IL-17A^+^ and Foxp3^+^ cells in PHB2-deficient T cells compared to control cells (Fig. 4B).

**Figure 4:**
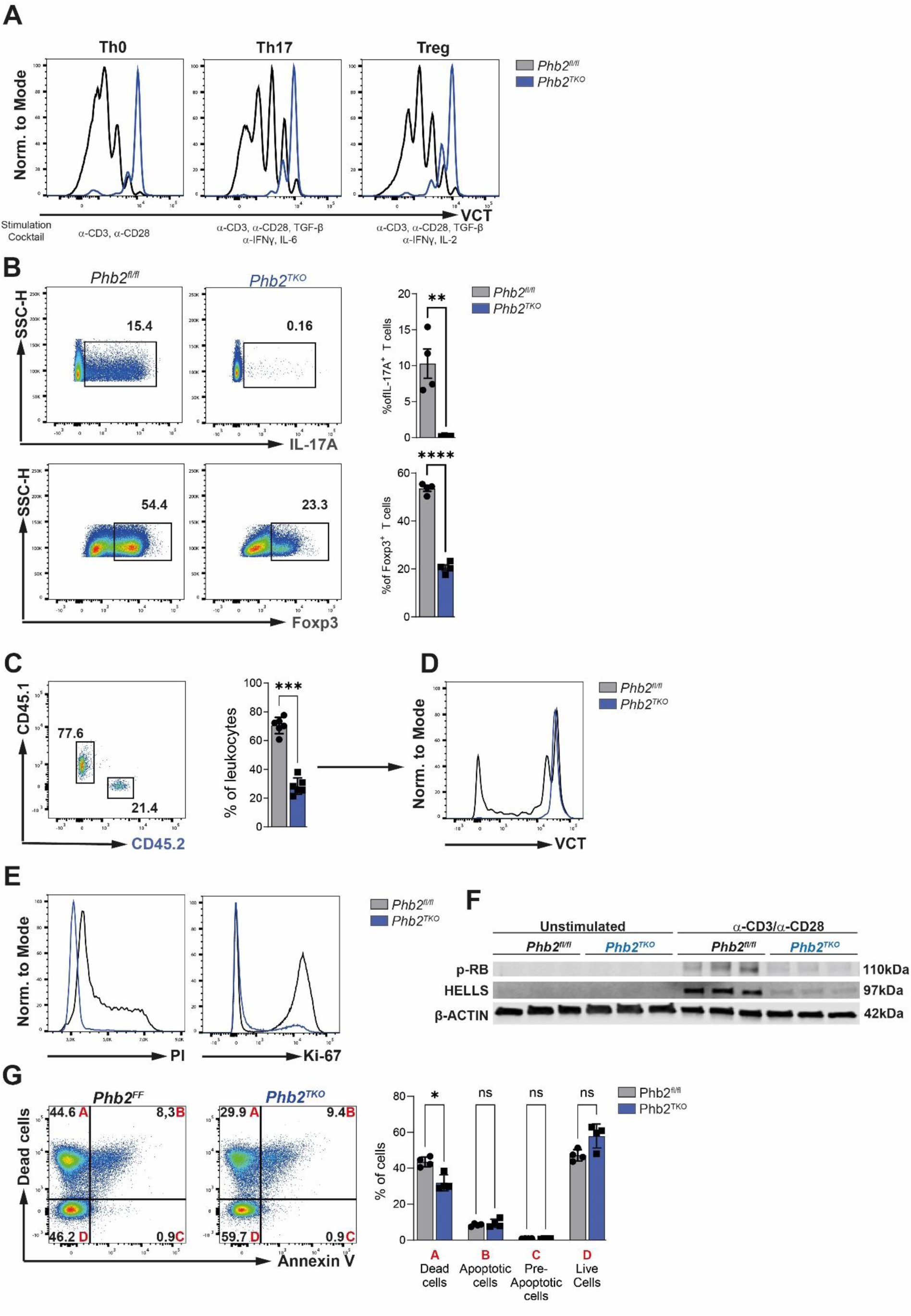
PHB2 deletion results in defects in T cell proliferation and differentiation. (A) Histogram of VCT on live CD4^+^ naïve T cells cultured under Th0, Th17 and Treg inducing conditions for 96h (n=4) (B) Flow cytometric (left) and statistical analysis (right) of IL-17A and Foxp3 expression of cells from A (n=4) (C) Flow cytometric (left) and statistical (right) analysis of the co-transfer of CD45.1 ^+^ control and CD45.2^+^ PHB2 deficient naïve CD4^+^ T cells into RAG 2^- / -^ mice (n=5) (D) Histogram of VCT on CD4 ^+^ T cells from B (n=5) (E) Histogram of cell cycle analysis (Propidium Iodide (PI)) and Ki-67 expression in CD4 ^+^ T cells that were stimulated in vitro for 24h with α-CD3/α-CD28 (n=3) (F) Western Blot analysis of p-Rb and HELLS expression in 24h unstimulated and 24h α-CD28/α-CD28 stimulated naïve CD4 ^+^ T cells (G) Flow cytometric (left) and statistical (right) analysis of apoptotic CD4 ^+^ T cells after 24h of in vitro stimulation with α-CD3/α- CD28 (n=4) Data are representative of three ( A-B, E-F) and one ( C-D) independent experiments. Analyses were performed on 8-12-week-old mice. Bar graphs show means +/- SDs and single values ( B, C, F) Statistical significance was calculated using unpaired two-tailed t-test with Holm-Šidák correction for multiple comparison. *p<0.05, **p≤0. 01, ***p≤0.001, ***p≤0.0001.

Since there was still a small proportion of T cells isolated from PHB2 deficient mice that was able to proliferate, as seen in Fig. 4A and Fig. S4A, we analyzed whether these cells possibly escaped CD4- cre mediated deletion of PHB2. Therefore, CFSE^low^ T cells (that strongly proliferated) and CFSE^high^ T cells (that did not proliferate) were FACS sorted for Western blot analysis of PHB2 expression. This analysis revealed that indeed, the few T cells that were able to proliferate still expressed PHB2 and escaped Cre-mediated recombination (Fig. S4A). Thus, PHB2 deficiency strongly suppresses T cell proliferation *in vitro*. These data are in line with previous reports, were PHB2 was proven to be essential for proliferation in other cells^1, 19^.

To evaluate whether the proliferative defect observed *in vitro* can also be translated to *in vivo* conditions, we co-transferred MACS sorted VCT labeled naïve CD4^+^ T cells with different congenic markers, (control T cells CD45.1^+^) and *Phb2^TKO^* T cells CD45.2^+^ in a ratio of 1:1 into *RAG2^-/-^* mice. After six days T cells in the spleens of *RAG2^-/-^* mice were analyzed by flow cytometry. Control CD45.1^+^ T cells made up 77,6% of all CD90.2+ T cells found in the spleens of *RAG2^-/-^* mice, while PHB2 deficient CD45.2+ T cells made up 21,4% (Fig. 4C). VCT staining showed that control CD45.1+ T cells successfully proliferated and started to fill the niche present in *RAG2^-/-^* mice (Fig. 4D), while CD45.2^+^ PHB2-deficient T cells did not proliferate (Fig. 4D).

In order to determine whether the significant reduction of peripheral T cells in PHB2 deficient mice is due to a block at a specific stage of T cell proliferation we performed a cell cycle analysis. Propidium iodide staining of naïve CD4^+^ T cells that were stimulated for three days with α-CD3/α-CD28 revealed a complete block of PHB2-deficient T cells to initiate DNA replication (Fig.4E, left). Furthermore, absent Ki-67 staining in PHB2-deficient T cells after three days of stimulation suggests a block in cell cycle progression at the G1 to S-phase transition, since Ki-67 is upregulated upon entry into the S-phase of cell cycle^20^ (Fig. 4E, right).

Our proteomics data (Figure 3) revealed that proliferation associated proteins are downregulated in stimulated PHB2-deficient T cells (Fig. 4C). Interestingly, one of the identified proteins HELLS together with UHRF1 was reported to be regulated by the Rb/E2f family contributing directly to retinoblastoma tumorigenesis^21^. Concomitantly, UHRF, which also establishes DNA methylation patterns, was also significantly downregulated in α-CD3/α-CD28 stimulated naïve CD4^+^ T cells compared to control cells (Fig. 3C, Table S1). Consequently, we asked the question whether the Rb/E2f axis, essential for the G1 to S-phase transition, is differential regulated in PHB2-deficient T cells. While HELLS protein was not detected in both unstimulated control and PHB2-deficient T cells, stimulation with α-CD3/α-CD28 revealed a robust upregulation of HELLS in control T cells. In contrast, naïve CD4^+^ T cells isolated from *Phb2^TKO^* were not able to upregulate HELLS expression after stimulation, indicating a role for PHB2 in the regulation of HELLS proteostasis (Fig. 4F). Additionally, western blot analysis revealed a lower phosphorylation for Rb protein in TCR stimulated PHB2 deficient naïve CD4^+^ T cells compared to control cells (Fig. 4D).

Another possible explanation for the decreased T cell numbers in *Phb2^TKO^* mice could be an increased sensitivity to apoptosis, leading to the demise of cells attempting to enter the S-phase of the cell cycle^22^. However, Annexin V staining of PHB2-deficient T cells stimulated for two days with α-CD3/α-CD28 revealed no difference in apoptotic cells, pre apoptotic cells or live cells compared to controls (Fig. 4F). Surprisingly, dead cells were significantly reduced in PHB2-deficient T cells compared to control cells (Fig. 4F).

In summary, PHB2-deficient T cells are unable to proliferate and subsequently differentiate into effector T cell subsets. Apoptosis as an explanation for reduced cell count and proliferation can be excluded since PHB2-deficient T cells did not exhibit increased sensitivity to apoptosis after stimulation which is in contrast to other PHB2-deficient types reported previously^1, 3^. Their failure to enter the S-phase of the cell cycle during proliferation and a reduction in RB phosphorylation suggests a general block of cell cycle progression at the G1- to S-phase transition, thus blocking proliferation and differentiation of activated T cells.

### PHB2 deficiency is indispensable for T cell activation

T cell activation, proliferation and differentiation into effector and memory T cells involve massive remodeling of T cell size, molecular content and create a massive increase in the demand for energy and amino acids^23^. In general, activation of T cells leads to an increase in cell size, also referred to as blasting^24^. The proliferative defect of PHB2-deficient T cells in response to TCR stimulation and T helper cell subset cytokine cocktails suggest possible alterations downstream of TCR signaling and its co-stimulatory receptor CD28. In order to test whether PHB2-deficient T cells are able to blast after TCR activation MACS purified naive CD4^+^ T cells were either left untreated or cultured for one or two days in the presence of α-CD3/α-CD28stimulation. FACS analysis revealed that PHB2-deficient T cells did not increase their size one day after stimulation and were less efficient in increasing their cell size compared to control T cells after two days of stimulation (Fig. 5A).

**Figure 5:**
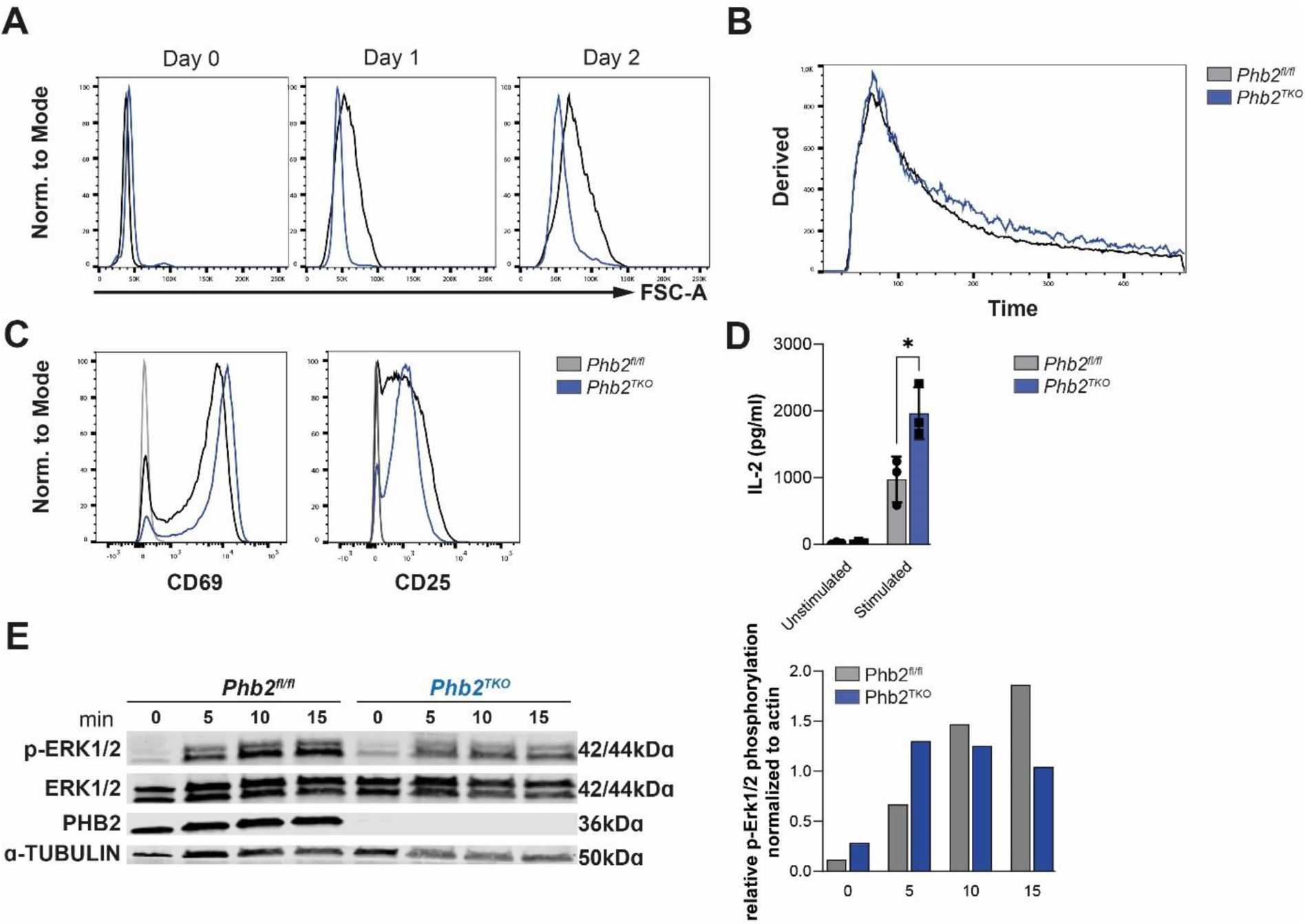
PHB2 deficiency does not influence the activation of T cells. (A) Histogram of cell size (FSC-A) of in vitro α-CD3/α-CD28 stimulated naïve CD4 ^+^ T cells for 0,1 and 2 days(n=4). (B) Calcium flux of naïve CD4 ^+^ T cells activated by biotin-α-CD3/CD4 cross-linkage (C) Histogram of CD69 and CD25 expression on naïve CD4 ^+^ T cells activated with α-CD3/α-CD28 for 24h. (n=4) (D) IL-2 secretion by naïve CD4 ^+^ T cells activated in vitro with α- CD3/ α- CD28 for 24h. (n=3) (E) Western blot analysis and normalization of Erk1/2 phosphorylation in CD4 ^+^ naïve T cells activated in vitro with α-CD3/α- CD28. (4 mice pooled) Data are representative of three (A-D) and one (E) independent experiment. Analyses were performed on 8-12-week-old mice. Bar graphs show means +/- SDs and single values ( D). Statistical significance was calculated using unpaired two-tailed t-test with Holm-Šidák correction for multiple comparison. *p<0.05.

The primary activation of T cells is mediated by the binding of the TCR to the MHCII molecule of antigen presenting cells. This receptor-ligand interaction leads to a sudden and very fast rush of Ca^2+^ into the cytoplasm mediated by calcium channels in the plasma membrane and the endoplasmic reticulum and subsequently promoting T cell proliferation^25^. We measured calcium flux by flow cytometry with the calcium sensitive dyes Fluo-4 and FuraRed. Calcium flux of PHB2 deficient naïve CD4^+^ T cells upon CD3 and CD4 cross-linkage revealed no significant difference in calcium influx compared to control T cells (Fig. 5B). Prolonged activation of T cells then leads to the upregulation of activation markers such as CD69 and CD25^26, 27^. Interestingly PHB2-deficient T cells showed no differences in CD69 and CD25 expression 24h after α-CD28/α-CD28 TCR stimulation compared to control T cells (Fig. 5C). Together this suggests that PHB2-deficient T cells retain functional signaling downstream of the TCR. The secondary activation through CD28 additionally induces the production and secretion of IL-2, a survival and pro-proliferative cytokine^28, 29^, which in turn can potentiate itself in a paracrine manner through IL-2 signaling via its receptor CD25^28^. We measured IL-2 secretion of naive PHB2 and control CD4+ T cells after 24h of α-CD28/α-CD28 stimulation by ELISA. Surprisingly, PHB2-deficient T cells produced significantly more IL-2 than control T cells (Fig. 5D).

Buehler et al. recently reported that plasma-membrane localized prohibitins affects Erk1/2 signaling thereby altering Th17 differentiation. To analyze Erk1/2 signaling of PHB2-deficient T cells we stimulated naïve CD4^+^ T cells with α-CD3/α-CD28 for 0, 5, 10 and 15 minutes. After 5 minutes of stimulation Erk1/2 phosphorylation was successfully upregulated in PHB2-deficient T cells compared to control T cells but declined at later timepoints while, as expected, the phosphorylation of control T cells further increased (Fig. 5E). However, as reported by Buehler et al., the inhibition of prohibitins on the plasma membrane, which solely reduces Erk1/2 signaling, did not lead to decreased proliferation. Therefore, our data does not suggest that this reduction in signaling alone can account for the lack of proliferation of PHB2- deficent T cells^10^.

These results, combined with the proteomics analysis, suggest that PHB2-deficient T cells are capable in the induction of their primary and secondary signals through the TCR and CD28 respectively.

### The mitochondrial structure and function of T cells are dependent on the presence of PHB2

Multiple studies have reported that the mitochondrial structure is disrupted in cells lacking PHB2^1, 3, 4, 30^. To assess the mitochondrial structure in PHB2-deficient T cells, we stained CD4^+^ T cells with Mitotracker Orange, a mitochondrial dye that is independent of the mitochondrial membrane potential. We found that, while control T cells possessed long concentrated tubular mitochondrial bodies, mitochondria of PHB2-deficient T cells were distributed in smaller bodies throughout the cell (Fig. 6A), displaying a fissioned phenotype, a process where mitochondria divide or segregate into two separate mitochondrial organelles. It was previously reported that the disruption of the homeostasis of the mitochondrial structure in PHB2 deficient cells is due to a disbalanced ratio of the two isoforms L-OPA1/S-OPA1^1^. Indeed, Western blot analysis of L-OPA1 of untreated as well as α-CD28/α-CD28 stimulated T cells showed a distorted ratio of S-OPA1/L-OPA1 in naïve CD4^+^ T cells isolated from *Phb2^TKO^* mice (Fig. 6B, C). To evaluate if this ultimately leads to dysfunctional mitochondria, we stained the mitochondrial membrane potential (ΔΨm) at steady state and after TCR stimulation with the potential dependent mitochondrial membrane dye TMRE. This analysis revealed a small but significant increase in membrane potential in PHB2-deficient T cells at steady state compared to control T cells (Fig. 6D). After stimulation, however, PHB2-deficient T cells did not increase their membrane potential compared to control T cells (Fig. 6D). Since T cells can increase their mitochondrial volume during activation up to fourfold^31^ and this could explain the difference in mitochondrial membrane potential after activation, we stained unstimulated and stimulated T cells with Mitotracker Green, a membrane potential independent mitochondrial dye, to investigate stimulation induced mitogenesis. Surprisingly, PHB2 deficient naïve CD4^+^ T cells displayed a similar increase in mitochondrial volume 24h after TCR stimulation compared to wildtype cells. (Fig. 6E). A dysfunctional membrane potential can be an indicator for a deficiency in the electron transport chain. Staining with CellROX, a ROS dependent dye, showed a reduced amount of ROS produced by PHB2-deficient T cells compared to control mice (Fig. 6F). Together, these findings suggests that PHB2-deficient T cells cannot increase ATP production upon activation since the ΔΨm directly drives the conversion of ADP to ATP by complex V of the ETC^32^.

**Figure 6:**
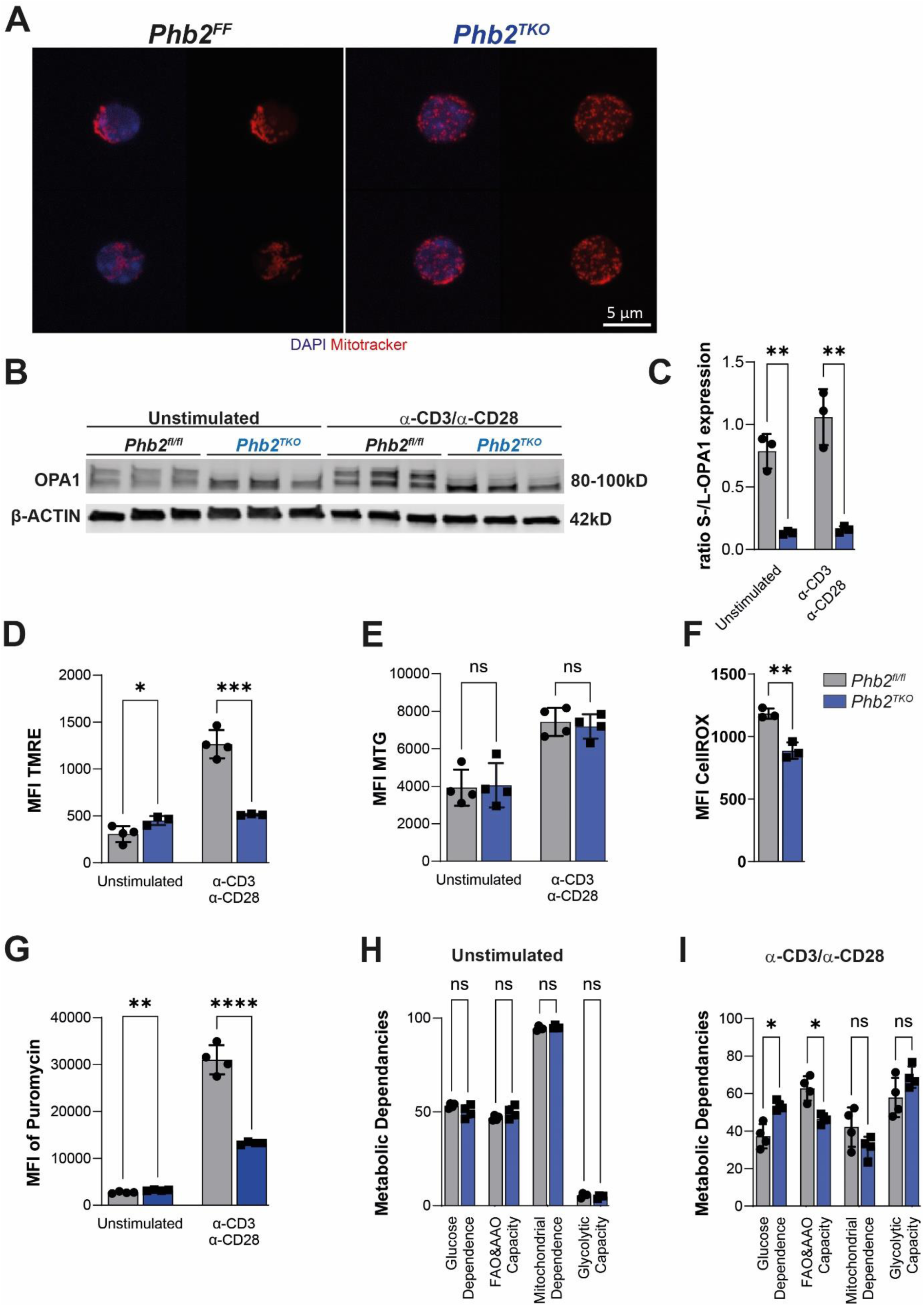
PHB2 is essential for mitochondrial structure and function. (A) Immunohistochemical staining of CD4+ T cells with DAPI and Mitotracker Orange CMTMRos. (B) Western Blot analysis of OPA1 expression in 24h unstimulated and 24h α-CD28/ α-CD28 stimulated naïve CD4 ^+^ T cells(n=3) (C) Statistical analysis of the relative S-/L-OPA1 ratio from B. (n=3) (D): The ΔΨm of naïve CD4 ^+^ T cells unstimulated or stimulated for 24h with α- CD28/ α-CD28. (n=3-4) (E) Mitochondrial mass of naïve CD4 ^+^ T cells unstimulated or stimulated for 24h with α-CD28/α- CD28 (n=4) (F) ROS in 24h α-CD28/α-CD28 stimulated CD4 ^+^ naïve T cells. (n=3) (G) SCENITH analysis of unstimulated and 24h a-CD28/a-CD28 stimulated CD4 ^+^ naïve T cells. Stimulated cells were pre-gated for CD25 (n=4) (H,I) Metabolic profile of unstimulated (H) and stimulated (I) CD4 ^+^ naïve T cells from G. Stimulated cells were pre-gated for CD25 (n=4) Data is representative of three (D, E, G-I), two ( B, C) and one (A, F) independent experiments. Analyses were performed on 8-12-week-old mice. Bar graphs show means +/- SDs and single values ( C-I). Statistical significance was calculated using unpaired two-tailed t-test with Holm-Šidák correction for multiple comparison. *p<0.05, **p≤0.01, ***p≤0.001, **** p≤0.0001.

To investigate if the reduced mitochondrial membrane potential seen after TCR stimulation translates into alterations in T cell metabolism, activated T cells were incubated with puromycin, an antibiotic that is incorporated into newly synthesized peptide chains at the ribosome during translation^33^. Since protein translation consumes up to 50% of the cell’s energy, any difference in energy generation also impacts *de novo* synthesis of proteins^34^. Analysis of activated T cells revealed that the expression of CD25 and puromycin incorporation correlated positively (Fig.S6A), thus we pre-gated on CD25 on stimulated T cells. At steady state PHB2-deficient T cells showed a small but significant increase in *de novo* protein synthesis. Interestingly, activated PHB2-deficient T cells upregulated protein translation after 24h of *in vitro* stimulation, however, did only upregulate protein synthesis to about half of that of control cells (Fig. 6G, Fig. S6B, C).

Next, naïve CD4^+^ T cells were treated with different metabolic inhibitors to analyze their metabolic profile by SCENITH^35^. Oligomycin was used to block mitochondrial derived ATP and 2-DG to block glycolytic derived ATP. The relative amount of protein synthesis that a cell can still conduct after treatment with one of these inhibitors indicates how dependent a cell solely is on glycolysis or mitochondrial ATP generation (Fig S6B, C). SCENITH analysis of naïve PHB2 deficient CD4^+^ T cells showed no differences compared to control cells (Fig. 6H). As reported previously for resting T cells, they mainly rely on mitochondrial respiration as their ATP source^36^. However, after α-CD28/α-CD28 stimulation, PHB2-deficient T cells showed a higher dependence on glycolysis, and a subsequent reduced capacity for fatty acid (FAO) and amino acid oxidation (AAO) (Fig. 6I).

Together this data show that PHB2-deficient T cells are unable to increase metabolic activity to meet the metabolic and anabolic demand after stimulation, thus possibly rendering them unable to proliferate.

## Discussion

Previous studies on PHB2 in T cells investigated T cell lines or targeted PHB2 specifically at the plasma membrane^10,10, 37^. By utilizing the CD4-cre mouse line we were able to investigate the role of PHB2 selectively in T cells *in vivo*. Analysis of these mice revealed a crucial role for PHB2 in the maintenance, differentiation, and activation of T cells. T cells were greatly reduced at steady state and unable to mount an effective immune response after immunization with MOG peptide. Furthermore, the genetic deletion of *PHB2* specifically in Treg cells led to a scurvy-like phenotype demonstrating an essential role for PHB2 in T reg cells.

We found, that although the thymic development was unaffected, all peripheral T cell populations were significantly reduced at steady state in spleen, LN and mLN in *Phb2^TKO^* mice. Interestingly, PHB2 deficiency did not manifest in the thymus of *Phb2^TKO^* mice, even though CD4-cre deletion is already active at the double positive stage during T cell development but was fully established in peripheral naïve T cells. One possible explanation could be that the PHB2 protein has a long half-life, and that cre mediated deletion only manifests in the periphery.

Some peripheral effector T cells still expressed PHB2, possibly due to the negative selection pressure and/or incomplete recombination by the cre. The fact that only PHB2 expressing T cells proliferated *in vitro* underscored this selection pressure imposed by PHB2 deficiency.

Previous studies on PHB2 in cell lines and mouse models already suggested that PHB2 has a conserved important role for different cell functions, such as proliferation and apoptosis^38–40^. We observed that PHB2-deficient T cells were incapable of proliferating. A proliferative defect has previously been reported in other PHB2-deficient tissues and cells^1, 3, 4^, and could be partially explained by the loss of L- OPA1, but it has not been specifically documented in T cells. These proliferation deficiencies were associated with increased sensitivity towards apoptosis in PHB2-deficient cells^1, 41, 42^. In one study on β-cells, the authors found that increased proliferation was accompanied by an increase in apoptosis^3^.

In contrast to other cell types, we did not detect an increased rate of apoptosis in stimulated PHB2- deficient T cells. Instead, cell cycle analysis revealed that the lack of proliferation may be due to a defect in entering the S phase and synthesizing DNA. This is further supported by the fact that we found S phase associated proteins in our proteomics data set, one of them being HELLS to be significantly downregulated. HELLS is a helicase that establishes DNA methylation patterns^43^ and has been shown to be essential for mature T cell proliferation^18^, thus representing one of the possible causes why PHB2- deficient T cells cannot proliferate.

In order to exclude the possibility that defective T cell receptor (TCR) activation is the cause for the proliferation defect, we performed calcium flux experiments upon TCR stimulation. Our results showed that PHB2-deficient T cells can initiate calcium flux. After 24 hours of stimulation, these cells also express the typical activation markers, such as CD25 and CD69. This suggests that signaling downstream of the TCR is intact, as CD69 expression is regulated by NF-κB, ERG-1, and AP-1^26^, while CD25 expression is regulated by NFAT and AP-1^44^.

Interestingly, PHB2-deficient T cells produced more IL-2 than control cells. IL-2 is induced by downstream TCR transcription factors (NFAT, AP-1, and NF-κB). This finding contradicts previous reports, which linked PHB2 to reduced IL-2 production in T cells. However, those studies focused on surface bound PHB2. This discrepancy suggests that surface bound and mitochondrial PHB2 perform distinct functions in T cells^45–47^. Another study showed that targeting surface localized prohibitins in T cells reduced MAPK-induced signaling, which reduced Th17 induction and promoted Treg cell induction, without affecting CD4 T cell proliferation^10^. Contrary to this, PHB2-deficient murine T cells exhibited a strong proliferation deficiency in our study.

A lack of PHB2 is associated with a fragmented mitochondrial phenotype, as documented in previous studies^1, 16^. This effect is directly linked to an imbalanced ratio between L-OPA1 to S-OPA1, resulting from the dysregulated processing of L-OPA1. In our study, we discovered that PHB2-deficient T cells also exhibit an imbalanced L-OPA1/S-OPA1 ratio and fragmented mitochondrial structure. Surprisingly, our proteomics data showed that the i-AAA protease YME1L1 is downregulated in activated T cells lacking PHB2. It has been previously demonstrated that YME1L regulates the processing of L-OPA1 through OMA1 and its absence leads to mitochondrial fragmentation^16, 48^. Furthermore, it was shown that it supports cell proliferation^48^. This may contribute to the defect in T cell proliferation; however, it does not account for the fragmentation of mitochondria at steady state.

Mitochondria of PHB2-deficient T cells displayed a normal membrane potential (ΔΨm) at steady state, but these cells failed to upregulate the ΔΨm upon TCR activation. This suggests a metabolic dysfunction, as the ΔΨm is crucial for the generation of ATP at complex V of the electron transport chain^32^. A subsequent analysis using the flow cytometry-based method to functionally profile metabolism with single cell resolution (SCENITH) revealed that T cells lacking PHB2 are incapable of enhancing their metabolic activity after activation. Specifically, we noticed a significant increase in glucose dependence and a decrease in fatty acid and amino acid oxidation of PHB2-deficient T cells compared to control cells. The disability to increases metabolic activity after activation implies that the proliferative defect of PHB2-deficient T cells may be due to their inability to support proliferation both at the metabolic and anabolic levels.

Surprisingly, our IPA analysis of stimulated PHB2-deficient T cells revealed no differentially regulated pathways related to T cell function or proliferation. However, the tRNA charging pathway was significantly upregulated in both unstimulated and stimulated PHB2-deficient T cells. This pathway involves aminoacyl-tRNA synthetases (aaRSs), enzymes that catalyze the esterification of amino acids to their corresponding tRNAs, a crucial step in the translation of proteins. There are two types of aaRSs: those in the mitochondria and those in the cytoplasm^17^. There was no bias between mitochondrial and cytoplasmic aaRS variants in our data, suggesting no preferential regulation of either type. The absence of a known master regulator of tRNA synthetases in T cells suggests a possible new direction for the function of PHB2.

In conclusion, this study shows that PHB2 is crucial for T cell homeostasis, proliferation, and effector function and possesses distinct functions in relation to their cellular location. Our findings show that PHB2 is indispensable for the proliferation and differentiation of T cells, with PHB2-deficient T cells exhibiting a significant cell cycle arrest during the transition from G1 to S phase. Moreover, PHB2- deficient T cells fail to upregulate their metabolism sufficiently after activation, resulting in diminished protein synthesis. These findings emphasize the importance of PHB2 in maintaining the integrity of T cell responses. Future research should investigate the molecular mechanisms underlying PHB2’s role in metabolic regulation and cell cycle progression, as well as its potential as a therapeutic target for immune-related disorders.

## Materials and Methods

### Mice

All mice used were on the C57BL/6J background and housed under specific pathogen-free conditions. *Phb2^fl/fl^* mice were crossed to CD4Cre mice to obtain *Phb2^TKO^* mice (B6-Phb2tm1Tlan Tg(Cd4-cre)1Cwi/Tarc).

### Single cell preparation

Single cell suspensions of thymus, spleen, lymph nodes and mesenteric lymph nodes were prepared by smashing organs through a 40µm filter in PBS supplemented with 2% fetal calf serum (FCS). Following, red blood cell lysis was conducted on spleenocytes.

### Antibodies and flow cytometry

Single cell suspension was used for flow cytometric stainings. First, Fc-block (5µg/mL) was conducted for 15 minutes at 4°C. Surface markers were then stained for 20 minutes at 4°C with antibodies against TCRb-Fitc (Biolegend #109205), TCRb-PE-Cy7 (Biolegend #109221), CD19-PE-Cy7 (Biolegend #115520), CD4-PerCP (Biolegend #100432), CD4-BV510 (Biolegend #100559), CD4-PE (Biolegend #100408), CD8-Pacifique Blue (Biolegend #100725), CD8-BV510 (Biolegend #100752), CD62L-APC (Biolegend #104412), CD44-PE (eBioscience #12-0441), CD44-Fitc (eBioscience #11-0441), CD25-Fitc (BD Biosciences #553072), CD45.1-PE-Cy7 (Biolegend #110730), CD45.2-Fitc (eBioscience #11- 0454), FVD-APC-eFl780 (eBioscience #65-0865), FVD-eFl506 (eBioscience #65-0866) and CD69-Fitc (eBioscience #11-0691). After staining cells were fixed in 2% PFA for 15 minutes at room temperature. For intracellular staining cells were stained for Ki67-APC (Biolegend #652405) and Foxp3-APC (eBioscience 17-5773) with the Bioscience™ Foxp3/Transcription Factor Staining Buffer Set (00-5523- 00).

Cells were acquired with a FACSCanto II cytometer (BD Bioscience) using FACS Diva software (BD Bioscience). Flow cytometry data was analyzed with FlowJo software version 10 (BD Bioscience). For all analyses, doublets (FSC and SSC properties) and dead cells (dye inclusion, fixable viability dye APC- ef780 eBioscience) were excluded.

### EAE Induction

Active EAE immunization with MOG_35–55_ (GenScript) peptide emulsified in complete Freund‘s adjuvant (CFA) (Difco) along with pertussis toxin (List Biological Laboratories) injections and disease assessment was performed as described elsewhere (REF). Brief, 50μg MOG35–55/CFA was injected subcutaneously into the tail base of the mouse. At the day of immunization and 2 days later, 150 ng pertussis toxin in PBS was applied i.p. unless otherwise stated.

### MACS purification and *in vitro* stimulation

CD4^+^ naïve T cells were purified using the naïve CD4+ T cell Isolation kit for mouse from Miltenyi Biotec (130-104-453). For short term kinetics 10×10^6^/ml naïve CD4 T cells were stimulated with 1µg/ml of α- CD3 and α-CD28 for the indicated time points. For 24h stimulations, 200 000/well naïve CD4 T cells were stimulated in a 96-flat well plate coated with 1µg/ml α-CD3 and added 1µg/ml α-CD28 to the medium.

For in vitro proliferation assay cells were labeled with CellTrace violet cell proliferation kit (VCT) according to the manufacturer (Invitrogen, C34557). For in vitro polarization, all cells were stimulated in a 96-flat well plate coated with 1µg/ml α-CD3 and added with 1µg/ml α-CD28 to the media. Additionally, 2ng/ml TGF-β, 25ng/ml IL-6 and 10µg/ml α-IFNγ for Th17 and 2ng/ml TGF-β, 10µg/ml α-IFNγ and 20ng/ml IL-2 for Treg polarization were added.

### Membrane Potential and MitoTracker Green staining

Stimulated naïve CD4 T cells were incubated with 25nM/ml TMRE (Abcam #AB113852) for membrane potential and 100nM Mitotracker Green FM (Thermo Fischer Scientific M7514) for mitochondrial mass for 40 minutes at 37°C and 5%CO_2_. Following cells were stained for surface markers as described above.

### Cell cycle Analysis

For cell cycle analysis cells were fixed with the eBioscience™ Foxp3/Transcription Factor Staining Buffer Set (00-5523-00). Next cells were incubated with RNAse (100µg(ml) for 15 minutes at 37°C. Propidium Iodide (50µg/ml, P4170 Sigma Aldrich) was added right before acquisition to the cells.

### Calcium flux

For calcium flux cells were stained with Fluo-4 (Thermo Fisher Scientific 1:250) and FuraRed (Thermo Fisher Scientific 1:100). For calcium flux induction cells were stained with 5µg/ml α-CD3 biotin (100304 Biolegend) and 5µg/ml α-CD4 biotin (13-0041 eBioscience). For cross connection 1mg/ml Neutravidin was added right before acquisition (Thermo Fischer Scientific 31000).

### SCENITH staining

For metabolic profiling, stimulated naïve CD4 T cells were incubated with metabolic inhibitors for 25 minutes. 2µM Oligomycin A (Sigma Aldrich 75351), 100mM 2DG (Sigma Aldrich D8375). Then, 10µg/ml Puromycin (Hycultec) was added for 40 Minutes. Cells were first stained for surface makers then stained with the anti-puromycin antibody (EMD Millipore MABE343-AF647) with the eBioscience™ Foxp3/Transcription Factor Staining Buffer Set (00-5523-00).

### Apoptosis essay

Stimulated naïve CD4 T cells were washed once with Annexin V staining buffer and then stained with Annexin V (Immunotools 31490016) antibody in Annexin V staining buffer for 20 minutes at 4°C. Annexin V staining Buffer 10x recipe: 0.1M HEPES, 1.4M NaCl, 25mM CaCl_2_.

### Western Blot

Cells were lysed in RIPA buffer (Cell Signaling Technology 9806) at 10×10^6^ cells per 50µl. PhosSTOP (Roche 04906837001) and protease inhibitor (Roche 04693132001) were added to the lysis buffer as per the manufacturer’s instructions. 10µg protein per sample was separated by gel electrophoresis on TGX Stain-Free Precast Gels (BioRad 4568083). PHB1 (Cell Signaling #2426) PHB2 (Biolegend #611801), b-actin (Cell Signaling #4970), HELLS (Cell Signaling #7998), Erk1/2 (Cell Signaling #9102), p-Erk1/2 (Cell Signaling #9106), a-tubulin (Cell Signaling #2144), OPA1 (BD #612606), p-rb (Cell Signaling #9309).

### Mitotracker Red staining

Cells were prepared at a concentration of 1×10˄6 cells/mL in 500 µL RPMI-1640 medium supplemented with heat-inactivated 10% FCS. After washing with RPMI-1640, 500 µL of MitoTracker Orange CMTMRos (M7510, Thermo scientific, 1 µM) was added, and cells were incubated at 37°C for 15 minutes. After washing the cells with 0.5% BSA/PBS, 500 µL of 4% PFA in PBS containing 0.2 µg/mL Hoechst 33342 was added and cells were incubated at room temperature for 10 minutes. Cells were washed twice with 0.5% BSA/PBS, resuspended in 50 µL of Mowiol 4-88 mounting medium, and mounted on glass slides with coverslips. The slides were incubated overnight before microscopy analysis.

### IL-2 ELISA

IL-2 in supernatant was measured with the BD OptEIA^TM^ Mouse IL-2 ELISA Set (BD Bioscience #555148) according to the manufacturer’s instructions.

### Mass Spectrometry

#### Proteomics sample preparation

Cells were lysed in 100 µL urea-containing buffer (7 M Urea, 2 M Thiourea, 1% Sigma Aldrich Phosphatase Inhibitor Cocktail 3 in 100 mM NH_4_HCO_3_) by sonication for 15 min (30 s on/off cycles) at 4 °C with high power in a Bioruptor device (Diagenode, Liège, Belgium). Subsequently, proteins were digested using filter aided sample preparation (FASP) as previously described in detail [1]. In brief, lysates were loaded onto spin filter columns (Nanosep centrifugal devices with Omega membrane, 30 kDa MWCO; Pall, Port Washington, NY) and washed three times with buffer containing 8 M urea. Afterwards, proteins were reduced and alkylated using DTT and iodoacetamide (IAA), respectively. After alkylation, excess IAA was quenched by the addition of DTT. Buffer was then exchanged washing the membrane three times with 50 mM NH_4_HCO_3_ and proteins digested overnight at 37°C using trypsin (Trypsin Gold, Promega, Madison, WI) at an enzyme-to-protein ratio of 1:50 (w/w).

After proteolytic digestion, peptides were recovered by centrifugation and two additional washes with 50 mM NH_4_HCO_3_. After combining peptide flow-throughs, samples were acidified with trifluoroacetic acid (TFA). A peptide aliquot corresponding to 5 µg of protein was lyophilized and reconstituted in 25 µL 0.1% formic acid (FA) for whole proteome analysis.

#### Liquid-chromatography mass spectrometry (LC-MS)

For the LC-MS analysis of the full proteome, 100 ng of the reconstituted peptides were separated on a nanoElute LC system (Bruker Corporation, Billerica, MA, USA) at 400 nL/min using a reversed phase C18 column (Aurora UHPLC emitter column, 25 cm x 75 µm 1.6 µm, IonOpticks) which was heated to 50°C. Peptides were loaded onto the column in direct injection mode at 600 bar. Mobile phase A was 0.1% FA (v/v) in water and mobile phase B 0.1% FA (v/v) in ACN. Peptides were separated running a linear gradient from 2% to 37% mobile phase B over 39 min. Afterwards, column was rinsed for 5 min at 95% B. Eluting peptides were analyzed in positive mode ESI-MS using parallel accumulation serial fragmentation (PASEF) enhanced data-independent acquisition mode (DIA) on a timsTOF Pro 2 mass spectrometer (Bruker Corporation) [2]. The dual TIMS (trapped ion mobility spectrometer) was operated at a fixed duty cycle close to 100% using equal accumulation and ramp times of 100 ms each spanning a mobility range from 1/K_0_ = 0.6 Vs cm^−2^ to 1.6 Vs cm^−2^. We defined 36 × 25 Th isolation windows from *m/z* 300 to 1,165 resulting in fifteen diaPASEF scans per acquisition cycle. The collision energy was ramped linearly as a function of the mobility from 59 eV at 1/K_0_ = 1.3 Vs cm^−2^ to 20 eV at 1/K_0_ = 0.85 Vs cm^−2^. All samples were measured in triplicates.

#### Raw data processing

Peptides were identified and label-free quantification (LFQ) of proteins was performed using DIA-NN (v1.8) [4]. Full proteome samples were processed using library free mode with standard parameters, except for tryptic cleavage sites considering no cleavage before proline. The FASTA protein database contained 17085 reviewed (Swissprot) protein entries of the mouse reference proteome and 172 common contaminant proteins and was obtained on 17.02.2022 from uniprot.org.

#### Statistical analysis

Common contaminants were filtered and proteins with at least 2 peptides were further considered. Fold change (FC) was calculated by median values for both conditions. If either of the conditions reported absence of intensities for all replicates, a fold change of 100 or 0.1 was assigned. Two sided t-test (assuming equal variances) was performed, if more than 60% of the replicates had quantitative values. If a protein was exclusively present in only one conditions, a p-value of 0.00001 was assigned. In case neither had more than 60% of values present across all replicates, a p-value of 1 was assigned. Benjamini-Hochberg (FDR) correction for multiple testing was applied on all p-values, including assigned values (1 and 0.00001). For the proteome dataset, proteins with FC ≥2 or ≤0.5 and adjusted p-value ≤0.05 were considered significant.

## Data availability

All raw files and processing results are available at https://repository.jpostdb.org/preview/122543010266eae68b46fa8 with the access key 1154.

## Statistics

All experiments undergoing statistical analysis included at least three or more biological replicates with number of repetitions as indicated in the figure legends. Statistical analyses were performed with Prism v9 software (GraphPad). Statistical tests were applied as indicated in the figure legends. All values are represented as mean ± SEM unless otherwise stated. *P* values were considered significant with p<0.05. No blinding of mice or samples was conducted. No statistical methods were used to determine the sample size.

## Study Approval

All animal experiments were approved by the local administrations (Landesuntersuchungsamt Koblenz, Germany). Experiments were performed in accordance with the guidelines from the translational animal research center (TARC) Mainz, Germany. All efforts were made to minimize suffering of the mice.

## Author contribution

JFV and CS performed experiments, analyzed data, prepared figures, and wrote the manuscript. PA, EZ performed experiments and analyzed data. TM, AN, UD and ST performed proteomics and bioinformatics analysis. TL provided mouse strains. SR, TW and AM were associated with conceptualization and supervision of the project and finalizing the manuscript.HY and KR helped with confocal imaging and western blot analysis. AW and NH supervised the project, acquired funding and wrote the manuscript together with JFV.

## Data Availability Statement

Text

## Materials & Correspondence

Dr. Nadine Hövelmeyer

Group leader

UNIVERSITÄTSMEDIZIN der Johannes Gutenberg-Universität

Institut für Molekulare Medizin

Geb. 308A, 1. OG, Zi. 1.203

Langenbeckstr. 1, 55131 Mainz

## Acknowledgments

We would like to thank Eva Schramm, Carsten Schelmbauer, Emmanouil Stylianakis and Cedric Bissinger for helping with the experiments.

