## Supplementary Information for "Prohibitin 2 is a Key Regulator of T Cell Proliferation and Effector Functions"

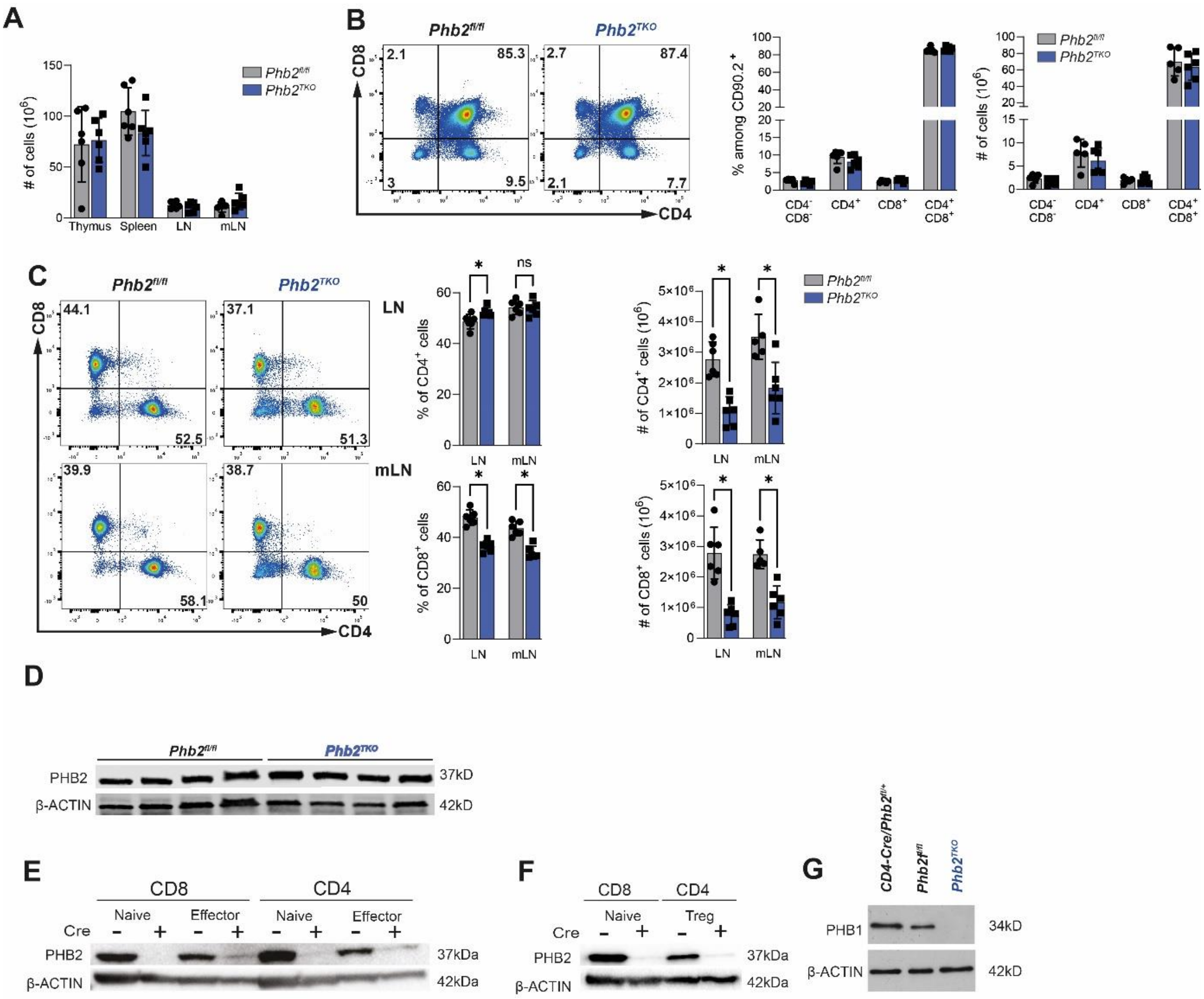

**Figure S1: Prohibitin 2 is essential for T cell homeostasis in vivo.**

(A) Total cell count of thymus, spleen, lymph nodes (LN) and mesenteric LN (mLN)

(B) Flow cytometric (left) and statistical (right) analysis of live CD90⁺ CD4⁺ and CD8⁺ thymocytes

(C) Flow cytometric and statistical analysis of live TCRβ⁺ CD4⁺ and CD8⁺ T cells in LN and mLN

(D) Western Blot analysis of PHB2 in MACS sorted CD4⁺ thymic T cells

(E) Western blot analysis of PHB2 in FACS sorted splenic naïve CD4⁺ (CD62L⁺ CD44^low), CD4⁺ effector (CD62L⁻ CD44^high), CD8⁺ naïve (CD62L⁺ CD44^low) and pooled CD8+ effector and central memory (CD62L⁻ CD44^high and CD62L⁺ CD44^high) T cells

(F) Western blot analysis of PHB2 of flow cytometry sorted splenic Treg cells (CD4⁺, CD25⁺), and naïve CD8⁺ (CD62L⁺ CD44^low) T cells

(G) Western Blot analysis of PHB1 in MACS sorted CD4⁺ splenic T cells

Data are representative of at least three independent experiments with n= 5 -6 mice per experiment per group. Analyses were performed on 8-12-week-old mice. Bar graphs show means +/- SDs and single values (A, C, F). Statistical significance was calculated using unpaired two-tailed t-test with Holm-Šidák correction for multiple comparison. *p<0,05.

.

**A**

**Unstim**

| | |
|---|---|
| PAXB1 | -6,64 |
| NUCG | -6,64 |
| PERE | -6,64 |
| HS71A | -6,64 |
| APC4 | -6,64 |
| TRFL | -6,64 |
| NTF2 | -6,64 |
| PPTC7 | -6,64 |
| UB2R2 | -6,64 |
| MTND | -6,64 |
| GILT | -6,64 |
| RM40 | -6,64 |
| ITIH2 | -1,57 |
| CD27 | -1,24 |
| COQ9 | -1,12 |
| EXOS4 | -1,06 |
| MOT1 | 1,02 |
| SYIC | 1,03 |
| SYNE3 | 1,05 |
| SBP1 | 1,08 |
| ATP5E | 1,16 |
| CHDH | 1,21 |
| TOP2A | 1,22 |
| SYEM | 1,31 |
| GLYM | 1,46 |
| GVIN1 | 1,49 |
| IGKC | 1,89 |
| IGHM | 2,57 |
| CHCH2 | 2,82 |
| PLCG2 | 6,64 |

**Shared proteins**

| | Unstim | Stim |
|---|---|---|
| PHB | -2,72 | -2,16 |
| PHB2 | -2,35 | -2,98 |
| DAPL1 | -2,30 | -6,64 |
| LETM1 | -1,41 | -1,34 |
| ABCB7 | -1,22 | -1,41 |
| GRP75 | 1,03 | 1,51 |
| CMPK2 | 1,18 | 1,69 |
| ANXA2 | 1,24 | 1,52 |
| SYYC | 1,34 | 1,27 |
| P5CS | 1,51 | 1,49 |
| 3HIDH | 1,54 | 1,37 |
| SYCC | 1,60 | 1,00 |
| S10A6 | 1,64 | 6,64 |
| SYAC | 1,96 | 1,38 |
| 4F2 | 2,09 | 1,11 |
| GRAA | 2,14 | 2,36 |
| GRAB | 2,31 | 6,64 |
| SCMC2 | 6,64 | 6,64 |
| TGM2 | 6,64 | 6,64 |
| MCPT8 | 6,64 | 6,64 |
| FCERG | 6,64 | 6,64 |

**Stim**

| | | | |
|---|---|---|---|
| UHRF1 | -1,67 | CYC | 2,11 |
| DNLI1 | -1,38 | S10A4 | 6,64 |
| NCBP2 | -1,26 | ANXA1 | 6,64 |
| MA7D1 | -1,19 | LEG1 | 6,64 |
| HELLS | -1,10 | MYO1F | 6,64 |
| YMEL1 | -1,02 | SPS2 | 6,64 |
| RCC2 | 1,04 | WBP2 | 6,64 |
| PCCB | 1,05 | ADSV | 6,64 |
| ZBP1 | 1,05 | DHSO | 6,64 |
| VIME | 1,06 | VINC | 6,64 |
| MTDC | 1,07 | PTN23 | 6,64 |
| IFIT1 | 1,08 | ELMO2 | 6,64 |
| STX7 | 1,08 | KS6A5 | 6,64 |
| VDAC2 | 1,09 | COR2A | 6,64 |
| SYSC | 1,09 | CLIC4 | 6,64 |
| ODPB | 1,15 | TOM1 | 6,64 |
| AB1IP | 1,15 | GSTM1 | 6,64 |
| STAT1 | 1,15 | PPOX | 6,64 |
| ROAA | 1,17 | VAMP3 | 6,64 |
| LAT1 | 1,19 | ASNS | 6,64 |
| IIGP1 | 1,19 | IRF9 | 6,64 |
| SFXN3 | 1,23 | THTM | 6,64 |
| SERA | 1,26 | GCC1 | 6,64 |
| SYWC | 1,39 | CHMP6 | 6,64 |
| 1433T | 1,44 | CATG | 6,64 |
| HA11 | 1,53 | BANP | 6,64 |
| GAPR1 | 1,53 | HB2A | 6,64 |
| NDRG1 | 1,54 | FSCN1 | 6,64 |
| K2C1 | 1,71 | DESP | 6,64 |
| | | PLAK | 6,64 |

Down log(FC) Up

**B**

**tRNA charging**

**Unstim**

| | |
|---|---|
| SYEM | 1,31 |
| SYIC | 1,03 |

**Shared proteins**

| | Unstim | Stim |
|---|---|---|
| SYAC | 1,96 | 1,38 |
| SYCC | 1,60 | 1,00 |
| SYYC | 1,34 | 1,27 |

**Stim**

| | |
|---|---|
| SYWC | 1,39 |
| SYSC | 1,09 |

**C**

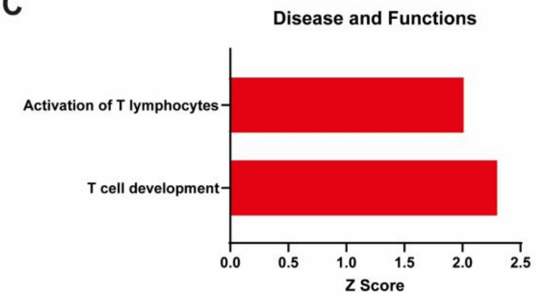


**Figure S3: PHB2 regulates proteins that are essential for proliferation.**

(A) Fold change list of significantly regulated peptides in unstimulated and α-CD3/α-CD28 stimulated control and PHB2 deficient naïve CD4$^+$ T cells

(B) Fold change of significant regulated tRNA transferase peptides

(C) Ingenuity Pathways Analysis of significantly regulated (Z Score ≥2) Diseases and Functions

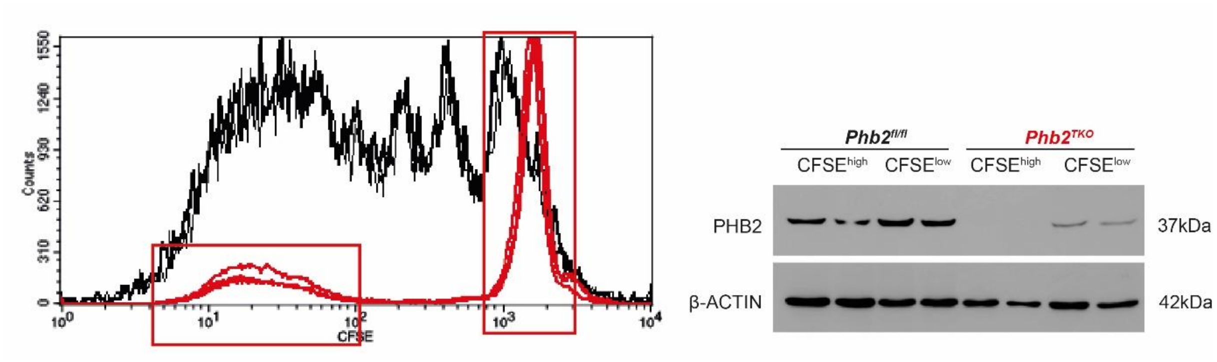


**Figure S4: PHB2 expressing T cells out proliferate PHB2 deficient T cells**.

Western blot analysis (right) of PHB2 in flow cyotmetric sorted (left) proliferating ($CFSE^{low}$) and non-proliferating ($CFSE^{high}$) $CD4^{+}$ T cells. Cells were stimulated in vitro for 4d with α-CD3/α-CD28. Data are representative of one experiment. (n=4). Analyses were performed on 8-12-week-old mice.

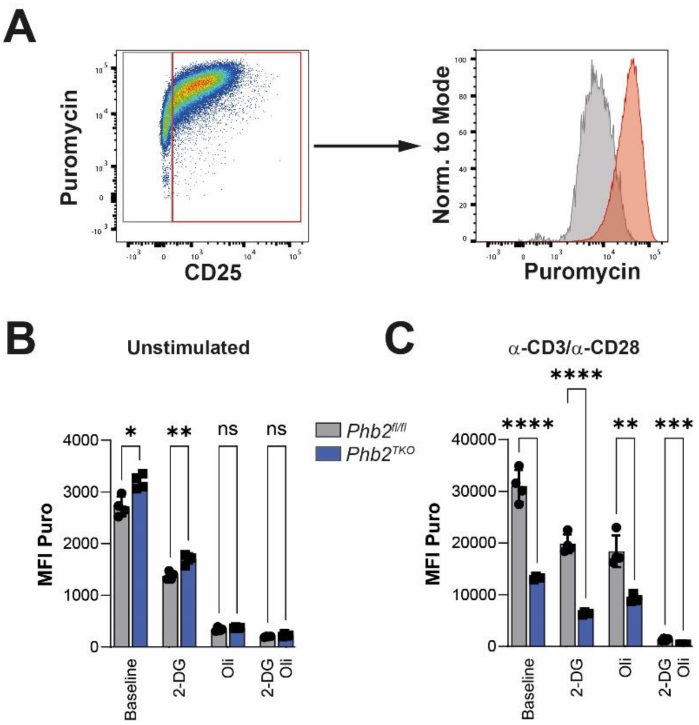


**Figure S6: CD25⁺ T cells incorporate more puromycin.**

(A) Flow cytometric analysis of Puromycin and CD25 co-expression (orange) on naive $CD4^+$ T cells stimulated for 24h with α-CD28/α-CD28 (n=4)

(B,C) MFI of Puromycin in 24h unstimulated (B) and 24 α-CD28/α-CD28 stimulated (C) $CD4^+$ naïve T cells (n=4)

Data is representative of three independent experiments. Analyses were performed on 8-12-week-old mice. Bar graphs show means +/- SDs and single values (C-I). Statistical significance was calculated using unpaired two-tailed t-test with Holm-Šidák correction for multiple comparison. *p<0.05, **p≤0.01, ***p≤0.001, ****p≤0.0001.

**Table S1: Proliferation associated proteins in PHB2 deficient T cells**

| Protein Name | Condition | Role described | Citations |
|---|---|---|---|
| HELLS | Stim | Essential for proliferation of peripheral T cells | T. M. Geiman and K. Muegge 2000 |
| DNLI1 | Stim | Immunodeficiency | P. Maffucci 2018, T. R. L. Howes and A.E. Tomkinson 2012 |
| UHRF1 | Stim | tumor progression, tumor severity, regulation of colonic T reg cell proliferation | S. C. Wu et al. 2022, Y. Obata et al. 2014 |
| RCC2 | Stim | progression in the metaphase | D. Papini et al. 2015 |
| NDRG1 | Stim | associated with T cell anergy induction | Y.M. Oh et al. 2015 |
| LEG1 | Stim | apoptosis and cell cycle bock at G1 | T. Lin et al. 2014 |
| BANP | Stim | activation of p53 and subsequent p21 | R. Kaul 2003, P. Bao 2023 |
| Paxbp1 | Unstim | Survival of T cells, checkpoint for cell growth, induction of p53 | W. Li 2023, S. Zhou 2021 |
| NCBP2 | Stim | expression positively correlates with tumor growth | H. Bu et al. 2022 |
